# Mobile genetic element-encoded hypertolerance to copper protects *Staphylococcus aureus* from killing by host phagocytes

**DOI:** 10.1101/279000

**Authors:** Marta Zapotoczna, Gus Pelicioli-Riboldi, Ahmed M Moustafa, Elizabeth Dickson, Apurva Narechania, Julie A. Morrissey, Paul J. Planet, Matthew T.G. Holden, Kevin J. Waldron, Joan A. Geoghegan

## Abstract

Pathogens are exposed to toxic levels of copper during infection and copper tolerance may be a general virulence mechanism used by bacteria to resist host defences. In support of this, inactivation of copper-exporter genes has been found to reduce the virulence of bacterial pathogens *in vivo*. Here we investigate the role of copper-hypertolerance in methicillin resistant *Staphylococcus aureus*. We show that a copper-hypertolerance locus (*copB-mco*), carried on a mobile genetic element, is prevalent in a collection of invasive *S. aureus* strains and more widely among clonal complex 22, 30 and 398 strains. The *copB* and *mco* genes encode a copper efflux pump and a multicopper oxidase, respectively. Isogenic mutants lacking *copB* or *mco* had impaired growth in subinhibitory concentrations of copper. Transfer of a *copB-mco* encoding plasmid to a naive clinical isolate resulted in a gain of copper hypertolerance and enhanced bacterial survival inside primed macrophages. The *copB* and *mco* genes were upregulated within infected macrophages and their expression was dependent on the copper sensitive operon repressor CsoR. Isogenic *copB* and *mco* mutants were impaired in their ability to persist intracellularly in macrophages and were less resistant to phagocytic killing in human blood than the parent strain. The importance of copper-regulated genes in resistance to phagocytic killing was further elaborated using mutants expressing a copper-insensitive variant of CsoR. Our findings suggest that the gain of mobile genetic elements carrying copper-hypertolerance genes contributes to the evolution of virulent strains of *S. aureus*, better equipped to resist killing by host immune cells.

## Introduction

Methicillin resistant *Staphylococcus aureus* (MRSA) is a major problem for animal and human health and is considered a global high priority pathogen by the World Health Organisation (1). One reason why MRSA continues to be a problem is that it evolves rapidly by acquiring mobile genetic elements (MGE) such as plasmids. Many successful contemporary clones of MRSA carry copper resistance genes located on MGEs (2-6) but the contribution of copper resistance to the fitness and virulence of *S. aureus* has not yet been studied.

Copper is a key component of innate immune bactericidal defences and macrophages use copper to kill intracellular bacteria by actively importing it into the phagosome (7-10). Eukaryotic copper transport is facilitated by CTR1-mediated import into the cell and ATP7a-dependent transport into the phagolysosome (7, 11). Under aerobic conditions, excess copper is proposed to catalyze the production of hydroxyl radicals via the Fenton and Haber-Weiss reactions, which may cause oxidative damage to macromolecules due to their high redox potential. Copper toxicity (under all or perhaps only anoxic conditions) involves the formation of adventitious Cu(I)-thiolate bonds, thus damaging enzymes that functionally require free cysteines or disulfide bonds, such as iron sulfur cluster proteins (12, 13). The toxic properties of copper are harnessed by host phagocytes, such as macrophages (11, 14). Infection signalling, which involves elevated levels of interferon gamma (IFNγ) and a release of copper into the plasma, may trigger activation of macrophages and increased import of copper, which enhances killing of phagocytosed bacteria (7, 10, 15).

Pathogens have evolved mechanisms to counteract copper toxicity, mainly by limiting the copper concentration in their cytoplasm through efflux or sequestration by copper metallochaperones, metallothioneins or storage proteins (16). Almost all bacteria possess genes that confer copper tolerance, from environmental bacteria isolated from black shale in copper-rich exploration regions (17) to human pathogens. Inactivation of copper-exporter genes has been shown *in vivo* to reduce the virulence of bacterial pathogens such as *Mycobacterium tuberculosis* (18), *Streptococcus pneumonia* (19), *Salmonella enterica (10)* and *Pseudomonas aeruginosa* (20). In some cases, the virulence defect has been shown to be due to the inability of these pathogens to resist copper-mediated killing within the macrophage phagosome (10). Data accumulated so far suggests that copper tolerance may be a general mechanism of virulence in bacteria and that pathogens are exposed to toxic levels of copper during infection (10, 18, 19, 21).

All *S. aureus* strains possess a conserved chromosomal operon, encoding the archetypal P_1_-type ATPase copper transporter CopA and a copper metallochaperone CopZ, that confers low level resistance to copper (Fig. 1A) (22). A copper-hypertolerance locus (*copB*-*mco*) has been described in some clinically relevant strains of *S. aureus*, carried either on a replicating plasmid or on a plasmid integrated into the chromosome (Fig. 1A) (2, 3, 5). The *copB* gene encodes a second copper exporting P1-type ATPase (CopB) and *mco* encodes a multicopper oxidase implicated in copper homeostasis and the oxidative stress response (23). A chromosomally-encoded homolog of the Cu-sensitive operon repressor (CsoR), first characterized in *M. tuberculosis* (24), was shown to control transcription of both operons in*S. aureus* (2).

**Figure 1.**
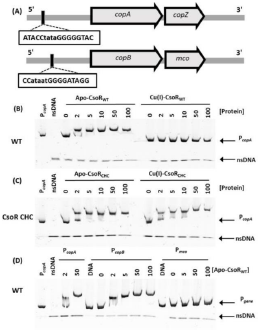
Electrophoretic mobility shift assay analysis of recombinant CsoR variants binding to the putative promoters upstream of *copA* (P*_copA_*), *copB* (P*_copB_*) and *mco* (P*_mco_*). A) Schematic representation of *copA*-*copZ* & *copB*-*mco* operons with putative promoter sequences. B-D) Recombinant wildtype CsoR_WT_ or the CsoR_CHC_ mutant (CsoR C41A/H66A/C70A) were purified and tested for binding to PCR products containing the DNA sequences (∼200 bp) upstream of the respective start codon, and a control DNA fragment of non-specific DNA sequence (nsDNA). B) Incubation of P*_copA_* DNA with wild type apo-CsoR, but not Cu(I)-CsoR, retards the migration of P*_copA_*. C) CsoR CHC retards migration of P*_copA_* in both the presence and absence of Cu(I). D) CsoR retards migration of P*_copA_* and P*_copB_* but not of*_mco_*

Here we investigated the role of copper-hypertolerance in *S. aureus*. We found that the MGE-encoded *copB* and *mco* genes improved bacterial growth under copper stress and enhanced bacterial survival within macrophages and in whole human blood. Expression of *copB* and *mco* was detected by intracellular bacteria isolated from macrophages and CsoR was responsible for regulating expression of these genes *in vivo*. Finally we determined the extent of carriage of *copB* and *mco* genes in a collection of invasive *S. aureus* isolates from European hospitals and in a more diverse collection of whole genome sequenced isolates from around the world.

## Results

### The tolerance of *S. aureus* to copper is enhanced by the *copB*-*mco* operons

The *copB*-*mco* copper-hypertolerance locus is carried either on a replicating plasmid or on a plasmid integrated into the chromosome (2, 3, 5). The role of MGE-encoded copper resistance genes in MRSA was studied using the *copB*-*mco* locus-carrying plasmid pSCBU, (3). Plasmid pSCBU, also known as P2-hm, was previously found to be carried by a population of MRSA CC22 bloodstream isolates from the UK and Ireland (3). For the purposes of this study, pSCBU was introduced into CC22 strain 14-2533T (Table S1). 14-2533T is a clinical isolate that is representative of the lineage where pSCBU was detected but it does not carry the plasmid. This strain was chosen as a clean and receptive host to study plasmid-conferred phenotypes.

The level of copper tolerance in 14-2533T carrying *copB* and *mco* genes on the replicating plasmid pSCBU was determined by measuring the minimal inhibitory concentrations (MICs) to copper salts (Table 1). Copper tolerance was the highest in 14-2533T carrying the replicating plasmid pSCBU (11 mM CuCl_2_), whereas the same strain without pSCBU had a lower MIC (6 mM). The individual contribution of *copB* and *mco* to copper tolerance was investigated by generating isogenic mutants carrying deletions in the copper tolerance genes on the plasmid pSCBU (pSCBUΔ*mco* and pSCBUΔ*copB*, Table S1 and Fig. S1). Deletion of *mco* or *copB* resulted in an MIC decrease to 8 mM or 6 mM CuCl_2_, respectively (Table 1), indicating these genes are the main contributors to pSCBU-mediated copper tolerance.

The pSCBU plasmid also encodes a cadmium-efflux system (*cadA*), which is known to protect from intracellular accumulation of toxic Cd(II), Zn(II) and Co(II) (25). As a control, cadmium and zinc tolerance of the pSCBU variants was tested. We observed that pSCBU conferred tolerance to cadmium and zinc (Table 1), which was unaffected by mutations in *copB* and *mco*, demonstrating that these genes do not influence resistance to these metals.

### CsoR binds to *copB* promoter DNA in a copper-dependent manner

The *S. aureus* copper-sensing transcriptional regulator (CsoR) was previously shown to negatively regulate both chromosomal and plasmid-encoded copper tolerance genes (2). Dissociation of CsoR from the GC-rich palindromic promoter regions has been shown to occur at two copper-regulated loci (*copA-copZ*, and *copB-mco*) in a copper-dependent manner (Fig 1A)(24). The *S. aureus* CsoR protein shares 24% amino acid sequence identity with CsoR from *Mycobacterium tuberculosis* (2). In *S. aureus*, residues Cys^41^, His^66^ and Cys^70^ are proposed to coordinate Cu(I) (26). Electrophoretic mobility shift assays (EMSA) assays performed anaerobically with recombinant CsoR and a ∼250 bp DNA fragment representing the *copA* promoter (P_*copA*_) confirmed that the wild type CsoR repressor bound specifically to the *copA* promoter, whereas anaerobic incubation with Cu(I) prevented association of CsoR with the promoter DNA (Fig. 1B). In contrast, a CsoR variant carrying C41A/H66A/C70A substitutions (CHC variant herein) remained bound to the *copA* promoter DNA despite the presence of copper (Fig. 1C), suggesting it is unable to coordinate Cu(I) and thus to undergo its copper-dependent allosteric conformational change. Thus the CsoR CHC variant is insensitive to copper, and de-repression of CsoR-regulated genes will not occur in cells expressing this variant.

Since *copB* and *mco* genes are responsible for hypertolerance to copper in *S. aureus* (Table 1), binding of CsoR to the *copB* and *mco* promoter regions was investigated. CsoR bound to DNA containing the sequence of the regions upstream of both *copA* and *copB* (P*_copB_*), but not that upstream of *mco* (P*_mco_*) (Fig. 1D), consistent with *copB* and *mco* being co-transcribed as part of an operon, under the regulatory control of CsoR through binding to P*_copB_*. There is no obvious CsoR binding sequence within the short intergenic region (14 bp) between *copB* and *mco* or in the 3’ sequence of *copB* (2).

**TABLE. 1.** Susceptibility of *S. aureus* strains to metals

| Strain | *copAZ* | *copB* | *mco* | MIC<sup>a</sup> Cu<sub>2</sub>SO<sub>4</sub> | *cadA* | MIC CdCl<sub>2</sub> | MIC ZnCl<sub>2</sub> |
| --- | --- | --- | --- | --- | --- | --- | --- |
| 14-2533T | ch<sup>b</sup> | NE<sup>c</sup> | NE | 6 mM | NE | 20 µM | 2 mM |
| 14-2533T (pSCBU) | ch | p<sup>d</sup> | p | 11 mM | p | 20 mM | 20 mM |
| 14-2533T pSCBUΔ*copB* | ch | Δ<sup>e</sup> | p | 6 mM | p | 20 mM | 20 mM |
| 14-2533T pSCBUΔ*mco* | ch | p | Δ | 8 mM | p | 20 mM | 20 mM |
| 14-2533T CHC | ch | NE | NE | 6 mM | NE | 20 µM | 2 mM |
| 14-2533T CHC pSCBU | ch | p | p | 10 mM | p | 20 mM | 20 mM |
| MRSA252 | ch | ch | ch | 8 mM | NE | NT<sup>f</sup> | NT |
| MRSA252 CHC | ch | ch | ch | 5 mM | NE | NT | NT |

<sup>a</sup>MIC, minimal inhibitory concentrations determined using a microdilution method, <sup>b</sup>ch, gene is incorporated into the chromosome, <sup>c</sup>NE, gene is not carried by the strain, <sup>d</sup>p, gene is carried on a replicating plasmid, <sup>e</sup>Δ, gene has been deleted by mutation, <sup>f</sup>NT, not tested.

### Copper-hypertolerance enhances growth of *S. aureus* at subinhibitory concentrations of copper

To study the role of the copper sensitive operon repressor CsoR in copper tolerance, site-directed mutagenesis was carried out on the *csoR* genes on the chromosomes of 14-2533T (CC22) and the CC30 strain MRSA252 to introduce amino-acid substitutions (C41A/H66A/C70A) that generate the copper-insensitive CsoR variant (CsoR CHC) (Table S1).

The MRSA252 strain carries a chromosomally-integrated plasmid carrying the *copB*-*mco* locus and was more tolerant to copper (MIC = 8 mM) than its isogenic CsoR CHC mutant (5 mM) (Table 1), showing that CsoR represses the copper-tolerance phenotype in MRSA252. In contrast, the CsoR CHC variant expressing strain 14-2533T CHC (pSCBU) had a similar MIC (10 mM) to the parent strain 14-2553T (pSCBU) (11 mM) and an elevated MIC compared to the plasmid-negative 14-2553T host strain (6 mM). This could reflect that CsoR does not fully repress *copB*-*mco* expressed from multi-copy plasmids, as shown previously by Baker *et al.* using a *csoR*-deficient mutant (2). Thus these data may indicate that the single-copy chromosomally–integrated *copB*-*mco* operon is more efficiently repressed by apo-CsoR than the pSCBU replicating plasmid-encoded operon.

To determine if expression of the *copB* and *mco* genes had an impact on bacterial growth under copper stress, we monitored the growth of cultures in TSB containing a concentration of copper below the MIC for all strains and mutants (4 mM, Fig. 2). Strain 14-2553T (pSCBU) grew faster and to a higher OD_600_ in sub-inhibitory concentrations of copper than the same strain without the plasmid or the mutants deficient in *copB* or *mco* (Fig. 2A). The defect in growth was more pronounced for the *copB* mutant than for the *mco* mutant. In contrast the 14-2533T CsoR CHC (pSCBU) mutant had an identical growth profile to the wild-type strain carrying the plasmid, again suggesting that CsoR does not fully repress *copB* and *mco* expression from the pSCBU plasmid (Fig. 2A). There was no growth advantage observed for strains growing in TSB lacking copper (Fig. 2B) or when low (μM) concentrations of copper salts were added to the growth medium (data not shown). As a control, the growth of MRSA252 and the MRSA252 CsoR CHC mutant were compared in TSB containing a sub-inhibitory concentration of copper (4 mM, Fig. 2C). MRSA252 grew more quickly and reached a higher OD_600_ than the copper-non-responsive regulatory mutant MRSA252 CsoR CHC, but not in media without copper (Fig. 2D). Transcription of *copB* and *mco* from pSCBU at sub-inhibitory concentrations of CuCl2 was quantified by Reverse Transcription quantitative PCR (RT-qPCR, Fig. 2E). A 5-to 12-fold increase in the abundance of transcript was measured for *mco* and *copB* in TSB cultures supplemented with sub-inhibitory concentrations of CuCl_2_ (4 mM), which confirmed that expression of *copB-mco* is copper-inducible. RNA transcripts of *mco* or *copB* were not detected in their respective deletion mutants (Fig. 2E, S1 & S3), as expected. Inducible copper-dependent expression of the other gene was detected in the mutants, showing that the respective gene deletions had not obstructed transcription of the other gene in this operon from the P*_copB_* promoter (Fig. 2E).

**Figure 2.**
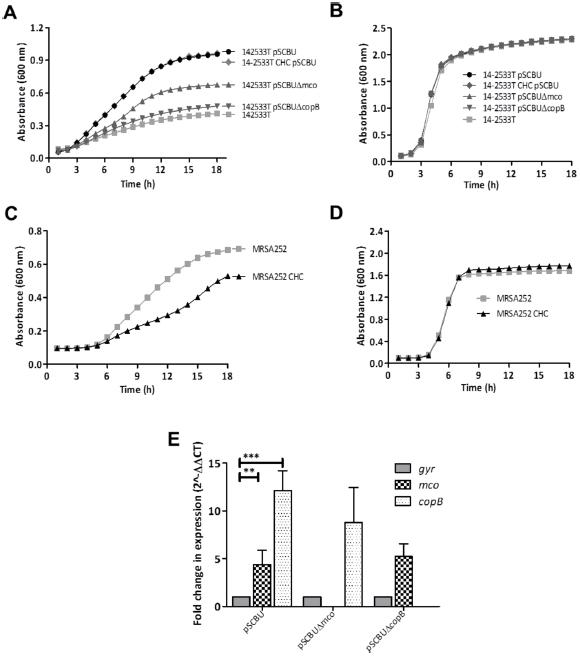
Enhanced growth in subinhibitory concentrations of copper chloride requires expression of copper tolerance genes. Growth of *S. aureus* 14-2533T and MRSA252 variants was measured in TSB supplemented with sub-inhibitory (4 mM) concentrations of copper chloride (A, C) or TSB broth alone (B, D). Growth curves representing data obtained from at least three independent experiments are presented. E) Fold change in expression of *copB* and *mco* in *S. aureus* cultured in TSB vs TSB with copper chloride (4 mM). The ΔΔCT method was used to determine the relative expression levels of the *copB* and *mco* genes of 14-2533T (pSCBU) and its mutants, normalised to *gyrB*. Presented are the means ± SD of 3 independent experiments, with statistical significance determined by ANOVA, *\*\*P<0.05 *** P<0.005.*

### Copper-hypertolerance genes increase *S. aureus* survival inside IFNγ-activated macrophages

Copper has previously been shown to be critical for the killing of bacteria following phagocytosis (7). In the presence of copper, activated macrophages up-regulate expression of the copper importer, CTR1, and commence trafficking of the P-type ATPase ATP7A to the phagolysosomal membrane, which leads to an enhanced killing of intracellular bacteria (7, 10).

To investigate whether bacterial tolerance to copper might influence the outcome for *S. aureus* following phagocytosis by macrophages, experiments were performed to quantify the survival of bacteria following phagocytosis. The murine macrophage cell line (RAW264.7) was activated with IFNɣ and treated with CuSO_4_ to induce expression of the relevant copper transporters (ATP7A and CTR1), which was confirmed using RT-qPCR (Fig. S2) (7, 27). IFNɣ-activated macrophages internalised the wild-type and mutants at similar levels (data not shown). However, 3 h post-phagocytosis, intracellular levels of bacteria were significantly different between the strains. The 14-2533T (pSCBU) strain survived inside the macrophages at significantly higher levels than 14-2533T without the plasmid (Fig. 3A). Importantly, the copper-susceptible *copB* and *mco* mutants had a survival defect compared to their parent strain 14-2533T, suggesting that copper tolerance in *S. aureus* prevents killing by macrophages (Fig. 3A). The CsoR CHC mutant of 14-2533T (pSCBU) did not show a significant survival defect in macrophages (Fig. 3A), probably reflective of the fact that plasmid-encoded CsoR-regulated genes are not efficiently repressed in this strain (Table 1, Fig. 2), thus it behaves like the wild-type. In contrast the MRSA252 CsoR CHC mutant had a defect in macrophage survival (Fig. 3B).

**Figure 3.**
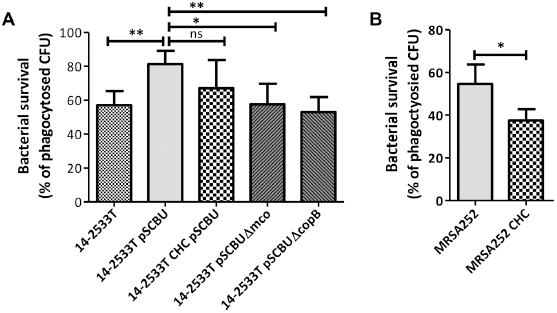
Hypertolerance to copper increases resistance of *S. aureus* to macrophage killing. Mouse macrophage cell line (RAW264.7) was suspended in DMEM supplemented with mouse IFNγ (40 ng/mL) and Cu2SO4 (40 μM) and seeded in 24 well plates at 2×106 cells per mL for 18h at 37°C in 5% CO2. 14-2533T (A) or MRSA252 (B) and derivatives were grown overnight in RPMI-1640 and then inoculated into the wells at an MOI of 10 in DMEM allowing phagocytosis for 30 min followed by killing of extracellular bacteria with gentamycin/lysostaphin for 30 min. Macrophages were then lysed at this time point (T0) and after 3h incubation (T3h) and subjected to viable count to determine the levels of bacterial survival. The CsoR C41A/H66A/C70A expressing mutants are indicated as CHC. Presented are means ± SD of three independent experiments. Statistical significance is indicated,**P<0.005 * P<0.05, ns, not significant.

To determine if the *copB* and *mco* genes are expressed by bacteria residing inside activated macrophages, RT-qPCR was performed using RNA obtained from intracellular bacteria at 3 h post infection. The relative transcription levels were compared between the wild-type strains and their isogenic CsoR CHC mutants. The *copB* and *mco* genes were found to be 44- and 28-fold upregulated, respectively, in wild-type MRSA252 compared to MRSA252 CHC recovered from infected macrophages (Fig. 4A). This demonstrated that a) *copB* and *mco* are expressed by *S. aureus* inside the macrophage, and b) that this expression is CsoR-dependent within immune cells (Fig. 4A). The same experiment was carried out with 14-2533T (pSCBU) and showed that *copB* and *mco* are expressed intracellularly in macrophages (Fig. 4B). However the increase in expression of *copB* and *mco* was much less for 14-2533T (pSCBU) than in the MRSA252 strain (Fig. 4B), which is consistent with susceptibility results (Table 1 and Fig. 2), indicating a weaker transcriptional control of CsoR over the plasmid-encoded genes compared to those that are carried on the chromosome of MRSA252.

**Figure 4.**
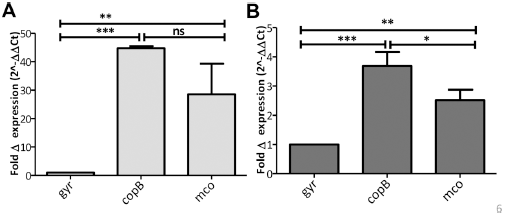
Intracellular expression of *copB* and *mco*. Fold change in expression of *copB* and *mco* by wild-type *S. aureus* relative to CsoR CHC mutants of either 14-2533T (pSCBU) (A) or MRSA252 (B). RAW264.7 macrophages were activated with IFNγ (50 µg/mL) and of copper chloride (40 µM) for 18 h. Infections of the macrophage monolayer were performed with RPMI-grown *S. aureus* at an MOI of 20. Extracellular bacteria were killed by treatment with with gentamycin/lysostaphin following washing of the monolayers with PBS. 3 h post infection RNA was isolated from infected macrophages and for to RT-qPCR. The ΔΔCT method was used to determine the relative expression levels of *mco* and *copB* in WT and CHC mutants normalised to *gyrB*. Presented are the means ±SD of 3 independent experiments, with statistical significance by ANOVA and Bonfferoni’s Multiple Comparison post test; *\*P< 0.5 **P<0.05 *** P<0.005.* ns, not significant

### Copper-hypertolerance genes increase survival of *S. aureus* in whole human blood

To determine whether the enhanced ability of *copB-mco* carrying strains to survive inside activated macrophages *in vitro* may be of relevance to infection of the human host, *ex vivo* infection studies were performed with whole human blood. Consistent with results obtained for intracellular survival within activated macrophages, copper-hypertolerant *S. aureus* 14-2533T (pSCBU) had an increased ability to survive in whole human blood compared to the 14-2533T strain without the pSCBU plasmid (Fig. 5A). This protection from killing in blood was due to copper resistance genes since the *mco* and *copB* mutants had a survival defect, similar to the plasmid-deficient 14-2533T strain (Fig. 5A). Protection from killing in blood could be attributed to resistance to phagocytic killing since incubation in the cell-free plasma fraction of the same blood under the same conditions yielded similar CFUs for wild-type and mutants (Fig. 5C).

**Figure 5.**
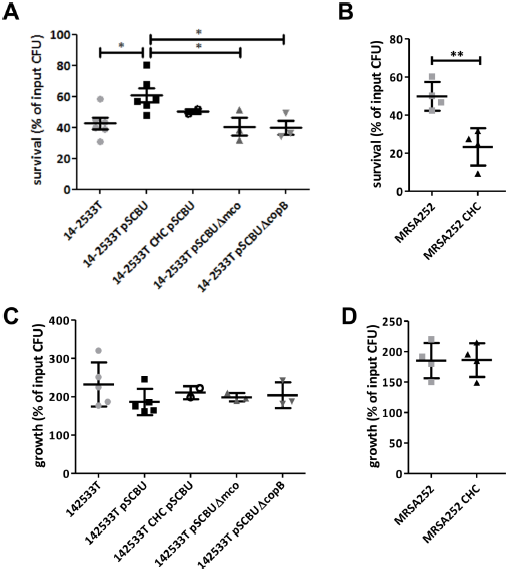
Increased survival of copper hypertolerant *S. aureus* in human blood. *S. aureus* (ca. 1×10^4^ CFU/mL) strains were inoculated into freshly drawn human blood (A, B) or plasma (C, D) and incubated for 3 h at 37°C. Viable count was used to determine the numbers of bacteria in blood or plasma. The number of CFU after 3 h is expressed as a percentage of the original input CFU at 0 h. Horizontal lines represent the means ± SD of at least three independent experiments. Statistical significance was determined using ANOVA following Dunnett’s Multiple Comparison Test, * P= 0.0192 (A) or an unpaired t-test, ** P= 0.0052 (B).

The CsoR CHC mutant of MRSA252 had a significant defect in whole blood survival compared to the wild type but did not show a defect in growth in plasma (Fig. 5B, 5D). This showed that failure to de-repress CsoR-regulated genes (Fig. 1) impaired the ability of *S. aureus* to survive in blood. Together these results show the importance of copper hypertolerance genes for *S. aureus* to resist cellular killing in human blood.

### The *copB-mco* operon is carried by invasive *S. aureus* isolates and by strains belonging to CC22, CC30 and CC398

The prevalence of the *copB-mco* operon was investigated by interrogating the whole genome sequences (WGS) of 308 invasive *S. aureus* isolates (28) from hospitals across Europe. Mapping the *copB*-*mco* sequences against the WGS showed that this operon was present in 55 of the invasive isolates (17.9%). The *copB* and *mco* genes were carried by isolates from two major clonal complexes (CC) within the population; CC22 and CC30, and also a single CC8 isolate. All CC30 strains carried the *copB*-*mco* operon. The most prevalent sequence type (ST) in the CC30 population carrying *copB*-*mco* was ST30, but ST2868, ST36 (EMRSA-16), ST2858, ST2864, ST2879, ST39, ST1829, ST2862, ST2881, and ST34 isolates also carried *copB*-*mco*. Among the CC22 strains, 50% were found to carry the operon and all of them belonged to ST22 apart from one ST2877 isolate. In summary, *copB*-*mco* was found to be present in invasive *S. aureus* strains from across Europe but predominantly in isolates from two important clonal groups, CC22 and CC30 (28).

To further explore the presence of *copB* homologs as well as related copper tolerance genes, we interrogated all publicly available *S. aureus* genomes (Genbank; n=8037). While a conserved *copA* was found universally in 99.9% of all genomes, *copB* homologs were the second most prevalent copper tolerance gene at ca. 34.4% of all the genomes. The *copB* and *mco* homologs were found mostly in CC22, CC30, and CC398, and only sporadically in other clonal complexes. To further characterize the distribution of genes in these three CCs, we constructed phylogenetic trees of each CC and mapped the presence and absence of each *copB* locus gene to each tree. Interestingly, the distributions of genes within each clade are strikingly different. For instance, CC30 genomes show a strong conservation of *copB* loci with very few predicted losses, whereas CC22 and CC398 have much more sporadic distributions that suggest multiple acquisitions and losses. This pattern could signal stronger, or more persistent, selection for *copB* loci in CC30 genomes as compared to CC22 and CC398, where selection may be weaker or intermittent. We also found evidence of a more diverse context to the copper hypertolerance genes than the original context in which they were found, i.e. the *copB*-*mco* operon, with an additional putative lipoprotein-encoding gene (4) (here called *copL*) frequently associated with the *copB*-*mco* operon in CC398 strains and less frequently in CC22 and CC30. These data indicate that the *copB*-*mco* copper hypertolerance genes are widely distributed in CC22, CC30 and CC398 and imply the presence of selection pressure for hypertolerance to copper.

## Discussion

The connection between gain of copper tolerance and increased virulence of several human pathogens has been reported over recent years. Here we demonstrate that *S. aureus* employs copper hypertolerance to resist macrophage killing and to survive in whole human blood. Presumably, better survival in human blood is due to an increased resistance to killing by the cellular component because control experiments indicated that growth in blood plasma was not affected by copper resistance genes (Fig. 2). The increased resistance to phagocytic killing conferred by the *copB*-*mco* locus is likely to impact on the virulence potential of the bacterium *in vivo* and may provide a selective advantage to the pathogen. Importantly, *copB* and *mco* were expressed within infected macrophages, and the expression of these genes was, at least partially, dependent on expression of copper-responsive CsoR (Fig. 4). This provides indirect evidence that the *copB*-*mco* operon is expressed intracellularly in macrophages, in response to copper.

By studying the genome sequences of a collection of invasive isolates obtained from hospitals across Europe we determined the prevalence of *copB-mco* to be 17.9% of all isolates, emphasizing the clinical relevance of this locus (5, 28, 29). The *copB* and *mco* genes were carried by all isolates belonging to CC30, by 50% of isolates from CC22 and by a single CC8 isolate. The plasmid carrying the *copB-mco* operon was also recently reported to be carried by 43-70% of bloodstream infection isolates of *S. aureus* (mostly CC22) from the UK and Ireland sampled between 2001 and 2010 (3). There is evidence of extensive loss and gain of the pSCBU (p2-hm) plasmid (3) highlighting the mobility of the *copB-mco* locus within populations of *S. aureus*. The global significance of copper resistance in *S. aureus* was further highlighted by the widespread presence of *copB* and *mco* in CC22, CC30 and CC398 strains (Fig. 6). Interestingly, CC398 are the most common CC found in European livestock. Previous studies have reported that 24.3% of livestock-associated MRSA carried the *copB* gene (30). The use of copper compounds as feed supplements in animal husbandry may be selecting for the carriage of copper resistance genes by MRSA (31). Copper hypertolerance in *S. aureus* is likely to have more broad implications for human health since the dominant clone of community associated (CA)-MRSA in North America (USA300) and a closely related CA-MRSA clone found in South America (USA300-LV) both independently acquired a putative copper resistance locus as part of the arginine catabolic mobile element and the copper and mercury resistance element, respectively (4). In both cases the copper resistance loci are adjacent to the SCC*mec* element.

**Figure 6.**
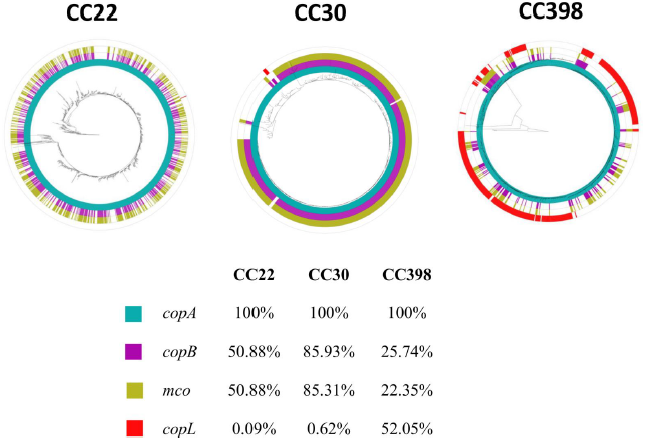
Maximum likelihood trees of CC22, CC30 and CC398 showing distribution of *copA, copB, mco, copL* genes. Trees are rooted in the longest branch (CC22 to *S. aureus* 08 01492; CC398 to *S. aureus* SO1977; CC30 to *S. aureus* MRSA252). Ring 1, 2, 3 and 4 show presence of *copA, copB, mco, copL* as blue, purple, yellow, and red.

Consistent with a previous report (2), our data show that the *copB-mco* operon mediates copper hypertolerance in *S. aureus*. Disruption of the *copB* or *mco* genes inhibited the growth of *S. aureus* in sub-inhibitory concentrations of copper demonstrating that carriage of both of these genes provides a fitness advantage to *S. aureus* under copper stress. It can be therefore concluded that both CopB-mediated copper efflux, and the activity of Mco, play a role in protecting *S. aureus* from copper.

The CsoR repressor, which has been previously implicated in transcriptional regulation of *copA*-*copZ* and *copB*-*mco* (2), was shown here to control expression of *copA* and *copB*-*mco* in a copper-dependent manner by binding directly to the DNA sequence upstream of *copA* and *copB* but not of *mco* (Fig. 1). Inactivation of the putative Cu(I)-coordinating residues Cys41, His66, and Cys70 (CHC) disrupted copper-dependent de-repression of CsoR-regulated genes, showing that *S. aureus* CsoR uses the same mechanisms of copper coordination as previously described by Liu *et al*. (24) for CsoR from *M. tuberculosis*. Although continued association of CsoR CHC with the *copA* and *copB* promoter DNA was confirmed by EMSA using recombinant proteins (Fig. 2), repression of the copper tolerance phenotype by the CsoR CHC variant was only completely effective in live bacteria with the chromosomally-encoded *copB*-*mco* operon (strain MRSA252 CHC). In contrast, the CHC mutant of 14-2533T (pSCBU) did not completely lose the copper hypertolerance phenotype shown by the parent strain carrying a wild-type copy of the *csoR* gene (Table 1, Fig. 2 and 4), showing that CsoR does not fully repress copper hypertolerance conferred by this plasmid. It may be that the pSCBU plasmid (are therefore *copB*-*mco*) are present in multiple copies, or due to poor diffusion of CsoR to the plasmid-located genes.

Horizontal gene transfer represents a major driving force in the evolution of *S. aureus* (32). This study provides important new insights into the contribution of MGE-encoded copper hypertolerance loci to the resistance of *S. aureus* to innate immune defenses. Due to the potential for MGEs to transmit rapidly in populations of *S. aureus*, our study shows that the spread of copper-hypertolerance loci could have important implications for the evolution of *S. aureus* as a pathogen.

## Materials and methods

### Bacterial strains and growth conditions

*S. aureus* strains used in the study are listed in Table S1. Bacteria were grown on tryptic soy agar (TSA) plates or in liquid cultures in ether tryptic soy broth (TSB) or RPMI-1640 at 37°C with shaking (200 rpm). To select for pSCBU-carrying strains, TSA was supplemented with CdCl_2_ at 1 mM. Growth curves were obtained using microtiter plates in TSB containing copper salts (either CuCl_2_ or CuSO_4_). For macrophage and whole blood survival assays bacterial strains were cultured in RPMI-1640 in aerated 50 mL falcon tubes at 37°C with shaking (200 rpm).

### Construction of mutations in plasmid-borne and chromosomally integrated *copB* &*mco*

Plasmid pSCBU was extracted from strain SASCBU26 (33) and used to transform strain 14-2533T (Table S1). Mutations in *S. aureus*, including deletions in the native plasmid pSCBU (Table S1; Fig. S1), were introduced using pIMAY (34). Plasmid deletions in the copper tolerance genes, pSCBUΔ*mco* and pSCBUΔ*copB*, were isolated in the 14-2533T (CC22) background (Table S1, Fig. S1). It was necessary to purify and reintroduce each validated mutated plasmid into a clean background in order to eliminate a mixed population containing a mutated and wild type copy of this multi-copy plasmid (2).

### Susceptibility testing

Minimal inhibitory concentrations (MICs) of soluble metal salts were determined using the standard broth microdilution method according to the guidelines by Clinical and Laboratory Standards Institute (CLSI). The lowest concentration of a compound showing no visible growth was recorded as the MIC.

### Production and purification of recombinant CsoR

The wild type *csoR* gene was amplified (Table S2) from *S. aureus* genomic DNA and cloned into pGEM-T (Promega). An internal NdeI site was mutated silently by QuikChange (Stratagene), and then *csoR* was sub-cloned into vector pET29a via NdeI/BamHI digestion and ligation. The *csoR* CHC mutant gene was amplified from the respective pIMAY construct. Constructs were confirmed by sequencing (GATC Biotech).

BL21(DE3) cells transformed with the resulting vectors, pET29a-CsoR or pET29a-CsoR-CHC, were cultured in lysogeny broth (LB) at 37°C with 180 rpm orbital shaking, and protein expression induced at OD_600_ ∼ 0.6 with addition of 1 mM isopropyl β-D-1-thiogalactopyranoside (IPTG), followed by further incubation at 30°C for 5 h. Cells were harvested, washed, resuspended in 25 mM Tris, pH 7.5, 15 mM dithiothreitol (DTT), containing protease inhibitor cocktail (Sigma) and lysed by sonication.

The supernatant was clarified by centrifugation and filtration, and purified by anion exchange chromatography on a 5 mL HiTrap Q HP column and an Akta purifier (GE Healthcare). Protein was eluted with a linear [NaCl] gradient (0-1 M NaCl), and CsoR-containing fractions (assessed by SDS-PAGE) subsequently concentrated on a 1 mL Heparin column (GE Healthcare) eluted with 1 M NaCl. This fraction was incubated overnight at 4°C with 10 mM EDTA and 20 mM tris(2-carboxyethyl)phosphine (TCEP), before resolution on a Superdex 75 16/600 column in 25 mM 4-(2-hydroxyethyl)-1-piperazineethanesulfonic acid (Hepes), pH 7.5, 200 mM NaCl, 15 mM DTT.

### EMSA

*S. aureus* MRSA252 genomic DNA was used to PCR amplify the putative promoter regions (i.e. the ∼200 bp upstream of the start codon) of *copA, copB*, and *mco* (Table S2), which were cloned into vector pGEM-T, confirmed by sequencing. The promoter fragments (plus ∼100 bp of flanking sequence from pGEM-T) were produced by PCR amplification from these pGEM-T constructs, plus a negative control fragment containing only the pGEM-T sequences. These PCR products were purified, and used in EMSA assays.

EMSAs were performed by incubating fully reduced (as determined with Elman’s reagent) recombinant CsoR variants (0-100 μM) with the respective promoter DNA plus the negative control DNA (both 0.1 μM) in 20 mM Hepes, pH 7.0, 100 mM NaCl, 100 ng/μ;L poly dI-dC (Sigma), 1 mM DTT, 0.4 mg/mL bovine serum albumin (BSA) at room temperature for 30 min. All incubations were performed anaerobically inside an N_2_ atmosphere glovebox ([O_2_] < 5 ppm – Belle Technology), and Cu(I)-CsoR was prepared by anaerobically incubating protein for 10 min with 1 mole equivalent of Cu(I) prepared as previously described (35). After incubation, samples were resolved on 6% acrylamide (w/v) native PAGE for 60-80 min at 82 V, and stained with 10% SYBR Safe solution (Invitrogen) for 20 min.

## RNA extraction

**To isolate RNA from *S. aureus***, bacterial cultures were grown in 20 mL TSB with or without copper salts (as indicated) to an OD_600_ ∼ 0,6. Cultures were suspended in phenol/ethanol (5:95%) mixture and incubated on ice for 1 h before pelleting the cells by centrifugation. At this step pellets were either stored at -70°C or subjected to total RNA extraction. To extract RNA, pellet(s) were gently suspended in 1 mL of TRIZOL following lysis using FastPrep^®^ Lysing beads (3×45 s, 2 min intervals on ice). Aqueous lysate was then mixed with chloroform (2:1) in Phase Lock Gel to separate the RNA-containing aqueous upper layer from the high density organic lower phase. The upper phase was precipitated with isopropanol (1:1) following ultracentrifugation at top speed for 30 min. The pellet was washed with 70% (v/v) ethanol and centrifuged. Supernatant was removed and the RNA pellet dried.

**RNA isolation from macrophages** was performed using a modified TRIZOL-based method. RAW264.7 cells were lysed directly in the culture dish by adding 12 mL of TRIZOL per T-175 cm^2^ flasks and scraping the cells. Chloroform was added to the suspension at 0.2 mL per 1 mM of TRIZOL reagent. Samples were immediately vortexed and incubated in RT for 2-3 min. Following centrifugation at 12,000 *g* for 15 min at 4 °C the mixture separated into layers and the upper aqueous layer was collected, precipitated with 0.5 mL isopropanol per 1mL of TRIZOL, incubated in RT for 10 min and centrifuged at 12,000 *g* for 10 min at 4°C. Obtained RNA pellet was washed once with 75% ethanol (adding at least 1mL per 1mL of TRIZOL).

**To isolate RNA from intracellular *S. aureus***, a combination of the above methods was used. Firstly, an infection was performed in T-175 cm^2^ flask following gentamycin/lysostaphin killing of extracellular bacteria and monolayer washing. Cells were then lysed with TRIZOL, like described above. Centrifugation at 4,000 *g* for 20 min was performed to separate the bacteria into a pellet. RNA from the bacteria-containing pellet and macrophage RNA –containing suspension were extracted using respective methods.

All air-dried pellets were dissolved in RNase-free molecular grade water and their stability and purity was checked by gel electrophoresis and the concentration determined using a Thermo Scientific Nanodrop spectrophotometer.

### RT-PCR

RNA was digested by DNase I treatment (Qiagen) according to manufacturer’s instructions and quantified using Nanodrop Spectrophotometer and the integrity assessed by electrophoresis. RNA was reverse transcribed to cDNA using High Capacity RNA-to-cDNA Kit (Applied Biosystems). RT-qPCR was performed using the Power SYBR Green PCR Master Mix (Applied Biosystems). The relative levels of gene expression in the treated cells and the non-treated controls were calculated by relative quantification using *gyrB* as the reference gene and using the primers in Table S2. All samples were amplified in triplicate and the data analysis was carried out using the Step One Software (Applied Biosystems).

Genomic DNA was isolated from cultured macrophages as described previously (http://cancer.ucsf.edu/_docs/cores/array/protocols/dna_cell_culture.pdf). Isolated DNA was used as a template to generate standard curve.

### Macrophage survival assays

Murine macrophage cell line (RAW264.7) was cultured in DMEM 10% (v/v) FBS. To generate monolayers 2×10^6^ cells per mL were seeded in in 24-well plates (500 µL per well) and incubated for 24 h in serum-free DMEM supplemented with CuSO_4_ (40 µM) and mouse IFNɣ (50 µg/mL) for 18 h at 37°C, 5% CO_2_. Immediately before the infection, RAW264.7 monolayers were washed with ice-cold DMEM alone. *S. aureus* strains were cultured in RPMI-1640. Immediately before the experiment bacteria were washed twice with DMEM and adjusted to an OD_600_ of 0.05 (ca. 2×10^7^ of CFU per mL) in DMEM and inoculated into the monolayers for 30 min. The monolayers were subsequently washed and extracellular bacteria were killed by treatment with gentamicin (200 μg/mL) & lysostaphin (100 μg/mL) for 30 min. Monolayers were then washed and lysed with ice-cold water at time point 0 (T0) and after additional 3 h incubation (T3) to determine the survival rates (CFU/mL). Lysates were plated on agar and CFUs were counted to determine numbers of viable bacteria.

### Human blood survival assays

The method for quantification of *S. aureus* in human blood was derived from Visai *et al.* (36). Briefly, *S. aureus* variants were grown in RPMI-1640 to stationary phase and diluted in RPMI and 25 µL (containing ca. 1×10^4^ CFU/mL) was added to 475 mL fresh blood obtained from healthy human volunteers that had been treated with 50 mg/mL of hirudin anticoagulant (Refludan, Pharmion). Tubes were incubated at 37°C with gentle rocking, and after 3 h serial dilutions were plated to determine the CFU/mL of viable bacteria. In parallel, an equal inoculum was incubated with cell-free plasma derived from the same donor’s blood. Bacterial numbers in plasma were quantified (CFU/mL) at 3 h time point and % survival of the original inoculum was determined. Ethical approval for the use of human blood was obtained from the TCD Faculty of Health Sciences ethics committee.

### Phylogenetic Matrix Construction and Gene Presence Absence

All preassembled genomes from public databases for CC22, CC30 and CC398 (n=1075, 320 and 707, respectively) were used for whole-genome alignment with reference to the *S. aureus* N315 genome, using the NUCmer and show-snps utilities of MUMmer (http://mummer.sourceforge.net) (37). The *S. aureus* genomes were assigned sequence types (ST) and CC by the *S. aureus* MLST typing scheme https://pubmlst.org/saureus/ sited at the University of Oxford (38) using the MLST typing perl script v. 2.9 for contigs (https://github.com/tseemann/mlst)and thereafter membership in each CC was determined by clade membership in a large (n=8037; unpublished data) *S. aureus* dataset composed of all publically available preassembled genomes. All regions from the reference genome annotated as mobile genetic elements were excluded. We also applied a mask that excluded repetitive sequences from the reference genome that were >80% identical over at least 100 nucleotides to other genomic loci, based on pairwise MegaBLAST-based analysis (39). For each CC, a Maximum likelihood phylogeny was constructed with RAxML v8.2.11 (40) using an ascertainment bias correction and the general time-reversible (GTR) substitution model (41) accounting for among-site rate heterogeneity using the Γ distribution and four rate categories (42) (ASC_GTRGAMMA model) for 100 individual searches with maximum parsimony random-addition starting trees. Node support was evaluated with 100 nonparametric bootstrap pseudoreplicates (43).

We used the *copA, copB* and *copL* genes from TCH1516 and *mco* from CA12 to search for closely related genes in the genus *Staphylococcus* in GenBank (wgs and nr databases, 9222 genomes as of 08/16/2017), using BLAST (tblastx with a cutoff value 1e-130 for *copA, copB* and *mco* while 1e-90 for *copL*) (44). The four genes were mapped to the three trees as highquality circular representations using GraPhlAn software tool (https://bitbucket.org/nsegata/graphlan/). The richness of the colour shows the percentage similarity with the seed sequence used.

### Statistics

The data presented by this study represent the means ± SD of three experiments unless otherwise stated. Statistical significance was assessed using two-way ANOVA and indicated as * for *P<0.05*, ** for *P< 0.001* and *** for *P< 0.0001*, unless otherwise stated.

## References

1. WHO. 2017. Prioritization of pathogens to guide discovery, research and development of new antibiotics for drug resistant bacterial infections, including tuberculosis World Health Organization.

2. Baker J, Sengupta M, Jayaswal RK, Morrissey JA. 2011. The *Staphylococcus aureus* CsoR regulates both chromosomal and plasmid-encoded copper resistance mechanisms. Environ Microbiol 13:2495–2507.

3. Jamrozy D, Coll F, Mather AE, Harris SR, Harrison EM, MacGowan A, Karas A, Elston T, Estee Torok M, Parkhill J, Peacock SJ. 2017. Evolution of mobile genetic element composition in an epidemic methicillin-resistant *Staphylococcus aureus*: temporal changes correlated with frequent loss and gain events. BMC Genomics 18:684.

4. Planet PJ, Diaz L, Kolokotronis SO, Narechania A, Reyes J, Xing G, Rincon S, Smith H, Panesso D, Ryan C, Smith DP, Guzman M, Zurita J, Sebra R, Deikus G, Nolan RL, Tenover FC, Weinstock GM, Robinson DA, Arias CA. 2015. Parallel Epidemics of Community-Associated Methicillin-Resistant *Staphylococcus aureus* USA300 Infection in North and South America. J Infect Dis 212:1874–1882.

5. Holden MT, Feil EJ, Lindsay JA, Peacock SJ, Day NP, Enright MC, Foster TJ, Moore CE, Hurst L, Atkin R, Barron A, Bason N, Bentley SD, Chillingworth C, Chillingworth T, Churcher C, Clark L, Corton C, Cronin A, Doggett J, Dowd L, Feltwell T, Hance Z, Harris B, Hauser H, Holroyd S, Jagels K, James KD, Lennard N, Line A, Mayes R, Moule S, Mungall K, Ormond D, Quail MA, Rabbinowitsch E, Rutherford K, Sanders M, Sharp S, Simmonds M, Stevens K, Whitehead S, Barrell BG, Spratt BG, Parkhill J. 2004. Complete genomes of two clinical *Staphylococcus aureus* strains: evidence for the rapid evolution of virulence and drug resistance. Proc Natl Acad Sci U S A 101:9786–9791.

6. Gomez-Sanz E, Kadlec K, Fessler AT, Zarazaga M, Torres C, Schwarz S. 2013. Novel erm(T)-carrying multiresistance plasmids from porcine and human isolates of methicillin-resistant Staphylococcus aureus ST398 that also harbor cadmium and copper resistance determinants. Antimicrob Agents Chemother 57:3275–3282.

7. White C, Lee J, Kambe T, Fritsche K, Petris MJ. 2009. A role for the ATP7A copper-transporting ATPase in macrophage bactericidal activity. J Biol Chem 284:33949–33956.

8. Hodgkinson V, Petris MJ. 2012. Copper homeostasis at the host-pathogen interface. J Biol Chem 287:13549–13555.

9. Djoko KY, Ong CL, Walker MJ, McEwan AG. 2015. The Role of Copper and Zinc Toxicity in Innate Immune Defense against Bacterial Pathogens. J Biol Chem 290:18954–18961.

10. Ladomersky E, Khan A, Shanbhag V, Cavet JS, Chan J, Weisman GA, Petris MJ. 2017. Host and Pathogen Copper-Transporting P-Type ATPases Function Antagonistically during *Salmonella* Infection. Infect Immun 85.

11. Festa RA, Thiele DJ. 2012. Copper at the front line of the host-pathogen battle. PLoS Pathog 8:e1002887.

12. Macomber L, Imlay JA. 2009. The iron-sulfur clusters of dehydratases are primary intracellular targets of copper toxicity. Proc Natl Acad Sci U S A 106:8344–8349.

13. Gaupp R, Ledala N, Somerville GA. 2012. Staphylococcal response to oxidative stress. Front Cell Infect Microbiol 2:33.

14. Achard ME, Stafford SL, Bokil NJ, Chartres J, Bernhardt PV, Schembri MA, Sweet MJ, McEwan AG. 2012. Copper redistribution in murine macrophages in response to *Salmonella* infection. Biochem J 444:51–57.

15. Hordyjewska A, Popiolek L, Kocot J. 2014. The many "faces" of copper in medicine and treatment. Biometals 27:611–621.

16. Gold B, Deng H, Bryk R, Vargas D, Eliezer D, Roberts J, Jiang X, Nathan C. 2008. Identification of a copper-binding metallothionein in pathogenic mycobacteria. Nat Chem Biol 4:609–616.

17. Dziewit L, Pyzik A, Szuplewska M, Matlakowska R, Mielnicki S, Wibberg D, Schluter A, Puhler A, Bartosik D. 2015. Diversity and role of plasmids in adaptation of bacteria inhabiting the Lubin copper mine in Poland, an environment rich in heavy metals. Front Microbiol 6:152.

18. Ward SK, Abomoelak B, Hoye EA, Steinberg H, Talaat AM. 2010. CtpV: a putative copper exporter required for full virulence of *Mycobacterium tuberculosis*. Mol Microbiol 77:1096–1110.

19. Johnson MD, Kehl-Fie TE, Klein R, Kelly J, Burnham C, Mann B, Rosch JW. 2015. Role of copper efflux in pneumococcal pathogenesis and resistance to macrophage-mediated immune clearance. Infect Immun 83:1684–1694.

20. Schwan WR, Warrener P, Keunz E, Stover CK, Folger KR. 2005. Mutations in the *cueA* gene encoding a copper homeostasis P-type ATPase reduce the pathogenicity of *Pseudomonas aeruginosa* in mice. Int J Med Microbiol 295:237–242.

21. Neyrolles O, Mintz E, Catty P. 2013. Zinc and copper toxicity in host defense against pathogens: *Mycobacterium tuberculosis* as a model example of an emerging paradigm. Front Cell Infect Microbiol 3:89.

22. Sitthisak S, Knutsson L, Webb JW, Jayaswal RK. 2007. Molecular characterization of the copper transport system in *Staphylococcus aureus*. Microbiology 153:4274–4283.

23. Sitthisak S, Howieson K, Amezola C, Jayaswal RK. 2005. Characterization of a multicopper oxidase gene from *Staphylococcus aureus*. Appl Environ Microbiol 71:5650–5653.

24. Liu T, Ramesh A, Ma Z, Ward SK, Zhang L, George GN, Talaat AM, Sacchettini JC, Giedroc DP. 2007. CsoR is a novel *Mycobacterium tuberculosis* copper-sensing transcriptional regulator. Nat Chem Biol 3:60–68.

25. Tynecka Z, Gos Z, Zajac J. 1981. Energy-dependent efflux of cadmium coded by a plasmid resistance determinant in *Staphylococcus aureus*. J Bacteriol 147:313–319.

26. Grossoehme N, Kehl-Fie TE, Ma Z, Adams KW, Cowart DM, Scott RA, Skaar EP, Giedroc DP. 2011. Control of copper resistance and inorganic sulfur metabolism by paralogous regulators in *Staphylococcus aureus*. J Biol Chem 286:13522–13531.

27. Watkins RL, Zurek OW, Pallister KB, Voyich JM. 2013. The SaeR/S two-component system induces interferon-gamma production in neutrophils during invasive *Staphylococcus aureus* infection. Microbes Infect 15:749–754.

28. Aanensen DM, Feil EJ, Holden MT, Dordel J, Yeats CA, Fedosejev A, Goater R, Castillo-Ramirez S, Corander J, Colijn C, Chlebowicz MA, Schouls L, Heck M, Pluister G, Ruimy R, Kahlmeter G, Ahman J, Matuschek E, Friedrich AW, Parkhill J, Bentley SD, Spratt BG, Grundmann H, European SRLWG. 2016. Whole-Genome Sequencing for Routine Pathogen Surveillance in Public Health: a Population Snapshot of Invasive *Staphylococcus aureus* in Europe. MBio 7.

29. Grundmann H, Aanensen DM, CC van den Wijngaard, Spratt BG, Harmsen D, Friedrich AW, European Staphylococcal Reference Laboratory Working G. 2010. Geographic distribution of *Staphylococcus aureus* causing invasive infections in Europe: a molecular-epidemiological analysis. PloS Med 7:e1000215.

30. Argudin MA, Butaye P. 2016. Dissemination of metal resistance genes among animal methicillin-resistant coagulase-negative Staphylococci. Res Vet Sci 105:192– 194.

31. Yazdankhah S, Rudi K, Bernhoft A. 2014. Zinc and copper in animal feed - development of resistance and co-resistance to antimicrobial agents in bacteria of animal origin. Microb Ecol Health Dis 25.

32. Lindsay JA. 2014. *Staphylococcus aureus* genomics and the impact of horizontal gene transfer. Int J Med Microbiol 304:103–109.

33. Harris SR, Cartwright EJ, Torok ME, Holden MT, Brown NM, Ogilvy-Stuart AL, Ellington MJ, Quail MA, Bentley SD, Parkhill J, Peacock SJ. 2013. Whole-genome sequencing for analysis of an outbreak of meticillin-resistant *Staphylococcus aureus*: a descriptive study. Lancet Infect Dis 13:130–136.

34. Monk IR, Shah IM, Xu M, Tan MW, Foster TJ. 2012. Transforming the untransformable: application of direct transformation to manipulate genetically *Staphylococcus aureus* and *Staphylococcus epidermidis*. MBio 3.

35. Vita N, Platsaki S, Basle A, Allen SJ, Paterson NG, Crombie AT, Murrell JC, Waldron KJ, Dennison C. 2015. A four-helix bundle stores copper for methane oxidation. Nature 525:140–143.

36. Visai L, Yanagisawa N, Josefsson E, Tarkowski A, Pezzali I, Rooijakkers SH, Foster TJ, Speziale P. 2009. Immune evasion by *Staphylococcus aureus* conferred by iron-regulated surface determinant protein IsdH. Microbiology 155:667–679.

37. Kurtz S, Phillippy A, Delcher AL, Smoot M, Shumway M, Antonescu C, Salzberg SL. 2004. Versatile and open software for comparing large genomes. Genome Biol 5:R12.

38. Jolley KA, Maiden MC. 2010. BIGSdb: Scalable analysis of bacterial genome variation at the population level. BMC Bioinformatics 11:595.

39. Morgulis A, Coulouris G, Raytselis Y, Madden TL, Agarwala R, Schaffer AA. 2008. Database indexing for production MegaBLAST searches. Bioinformatics 24:1757–1764.

40. Stamatakis A. 2014. RAxML version 8: a tool for phylogenetic analysis and post-analysis of large phylogenies. Bioinformatics 30:1312–1313.

41. Lanave C, Preparata G, Saccone C, Serio G. 1984. A new method for calculating evolutionary substitution rates. J Mol Evol 20:86–93.

42. Yang Z. 1994. Maximum likelihood phylogenetic estimation from DNA sequences with variable rates over sites: approximate methods. J Mol Evol 39:306–314.

43. Felsenstein J. 1985. Confidence Limits on Phylogenies: An Approach Using the Bootstrap. Evolution 39:783–791.

44. Altschul SF, Gish W, Miller W, Myers EW, Lipman DJ. 1990. Basic local alignment search tool. J Mol Biol 215:403–410.

